# Identification of Differentially Expressed Gene Modules in Heterogeneous Diseases

**DOI:** 10.1101/2020.04.23.055004

**Authors:** Olga Zolotareva, Sahand Khakabimamaghani, Olga I. Isaeva, Zoe Chervontseva, Alexey Savchik, Martin Ester

## Abstract

**Motivation:** Identification of differentially expressed genes is necessary for unraveling disease pathogenesis. This task is complicated by the fact that many diseases are heterogeneous at the molecular level and samples representing distinct disease subtypes may demonstrate different patterns of dysregulation. Biclustering methods are capable of identifying genes that follow a similar expression pattern only in a subset of samples and hence can consider disease heterogeneity. However, identifying biologically significant and reproducible sets of genes and samples remains challenging for the existing tools. Many recent studies have shown that the integration of gene expression and protein interaction data improves the robustness of prediction and classification and advances biomarker discovery.

**Results:** Here we present DESMOND, a new method for identification of Differentially ExpreSsed gene MOdules iN Diseases. DESMOND performs network-constrained biclustering on gene expression data and identifies gene modules — connected sets of genes up- or down-regulated in subsets of samples. We applied DESMOND on expression profiles of samples from two large breast cancer cohorts and have shown that the capability of DESMOND to incorporate protein interactions allows identifying the biologically meaningful gene and sample subsets and improves the reproducibility of the results.

**Availability:** https://github.com/ozolotareva/DESMOND

**Supplementary information:** Supplementary data are available at *Bioinformatics* online.

## 1 Introduction

Since the development of any disease is thought to be due to dysregulation of certain molecular processes, the detection of genes differentially expressed in disease is necessary for understanding its pathogenesis. Many existing methods (McCarthy *et al.*, 2012; Love *et al.*, 2014; Ritchie *et al.*, 2015) are aimed at detecting genes which are significantly differentially expressed in the disease group compared to the control group. However, the reproducibility of differentially expressed gene discoveries is rather low (Zhang *et al.*, 2008; Sweeney *et al.*, 2016) because of high levels of noise and large technical variation. One possible way to improve reproducibility and consistency between dysregulated genes detected in independent studies is to group individual genes demonstrating a similar pattern of dysregulation together. Aggregating expressions of multiple genes into single values reduces the dimensionality of the data and facilitates the subsequent analysis. The resulting groups of coordinately expressed genes, also called *gene modules* (Saelens *et al.*, 2018; Mitra *et al.*, 2013), are likely to be functionally related and easier to interpret than the whole list of differentially expressed genes. These gene modules may be known *apriori*, or detected from the data when gene expression profiles are mapped on the interaction network. For example, active subnetwork detection methods search for sets of differentially expressed genes connected in the interaction network. These methods search for active subnetworks in a supervised manner, when class labels, e.g. disease and control or disease subtypes, are known (Ideker *et al.*, 2002; Chowdhury and Koyutürk, 2009; Dao *et al.*, 2011). However, class labels are not always available, and even if they are provided, some classes may be internally heterogeneous and consist of several latent molecular subtypes (McClellan and King, 2010; Perou *et al.*, 2000).

Biclustering methods (Pontes *et al.*, 2015; Padilha and Campello, 2017; Xie *et al.*, 2018) perform an unsupervised search for subsets of genes demonstrating similar expression patterns in a subset of samples, given a matrix of genes profiled in these samples. Since biclustering is a much more complex problem than clustering due to the much larger size of the search space, many biclustering methods put additional constraints on the input data or the biclustering result, e.g. they assume a hidden checkerboard data structure (Turner *et al.*, 2005; Hochreiter *et al.*, 2010) or binary expression values (Prelić *et al.*, 2006; Serin and Vingron, 2011). Although most of the biclustering methods work on expression or other omics data matrices (Khakabimamaghani and Ester, 2015), some of them can additionally incorporate orthogonal biological data to improve biclustering results. For example, COALESCE can optionally accept sequences and perform *de novo* motif search jointly with biclustering (Huttenhower *et al.*, 2009). It searches for biclusters composed of genes whose regulatory regions are enriched by the same motifs. Another biclustering method, cMonkey2, in addition to motif enrichment data, considers functional associations between genes in its scoring function (Reiss *et al.*, 2015). As well as many other biclustering methods, cMonkey2 is more suitable for the detection of differentially co-expressed biclusters, rather than differentially expressed. The new version of QUBIC can utilize the data on known gene relationships when ranks gene pairs before constructing biclusters (Li *et al.*, 2009).

Indeed, taking into account the known interactions between the genes may reduce the complexity of the problem. Instead of considering all possible gene subsets, in network-constrained biclustering, the solution is searched among interacting genes, which guides the method to more biologically reliable results. These considerations motivated us to develop DESMOND, a new method for identification of **D**ifferentially **E**xpre**S**sed gene **MO**dules i**N D**iseases. The advantage of DESMOND in comparison with most existing biclustering methods is its ability to incorporate prior knowledge about gene interactions which promises to improve the quality of the results. Another advantage of DESMOND is that instead of setting a hard binarization threshold it applies flexible thresholds for identification of samples in which genes are differentially expressed.

We conducted experiments on synthetic data and on real expressions from two large breast cancer cohorts. Breast cancer was chosen as an example of a heterogeneous disease with multiple characterized molecular subtypes. Various classifications of breast tumors that take into account various tumor characteristics and result in different numbers of breast cancer subtypes include the classification based on ER, PR and Her2 expression (Gradishar *et al.*, 2017), intrinsic subtypes classification (Perou *et al.*, 2000), WHO histopathological classification (for Research on Cancer, 2012), etc. We compared DESMOND with nine state-of-the-art biclustering methods (Cheng and Church, 2000; Lazzeroni and Owen, 2000; Murali and Kasif, 2003; Bergmann *et al.*, 2003; Li *et al.*, 2009; Huttenhower *et al.*, 2009; Hochreiter *et al.*, 2010; Serin and Vingron, 2011; Rodriguez-Baena *et al.*, 2011) (Supplementary Table 1) chosen based on their good performance on synthetic datasets with differentially expressed biclusters (Padilha and Campello, 2017). Among those, only QUBIC was able to take into account network data.

## 2 Methods

### 2.1 Problem definition

The problem addressed in this paper is the discovery of connected groups of genes differentially expressed in an unknown subgroup of samples, given a network of gene interactions and a matrix of gene expression profiles. This problem can be classified as network-constrained biclustering, or, alternatively, as unsupervised active subnetwork detection, when the desired sample subgroups are unknown.

Formally speaking, given expressions of genes in *G* measured in the samples of set *S*, and an undirected and unweighted graph *N* = (*G, I*), representing *I* interactions between the *G* genes, we aimed to find subsets of *G′* ⊂ *G* genes and *S′* ⊂ *S* samples, such that genes *G′* are differentially expressed in a subset of samples *S′* compared to the background samples 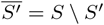; and *G′* forms a connected component in the network *N*. We call such pairs (*G′*, *S′*) modules. We define gene *g* to be differentially expressed in a set of samples *S′* ⊂ *S* compared to 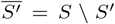, if *µ*_*g,S′*_, its median expression in *S′*, is different from the median expression 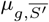 in 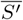. Since we are interested in discovering subgroups of genes that differentiate disease subtypes and may be used as biomarkers, in this work we employ the signal-to-noise ratio (SNR) (He and Zhou, 2008; Mishra and Sahu, 2011) as a measure of differential expression. The SNR for expression of gene *g* in *S′* samples is defined as

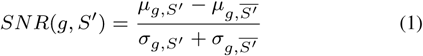

Similarly, we call a set of genes *G′* differentially expressed in the samples of set *S′* if ∀ gene *g* ∈ *G′*, *g* is differentially expressed in *S′*. As a measure of differential expression of a bicluster *B*(*G′*, *S′*), we use the average of absolute SNR over all genes *G′* in *S′* samples. Higher average absolute SNR indicates that a subset of samples *S′* is well-separated from the background in a subspace of *G′*. Such gene sets are promising biomarker candidates for distinguishing unknown but biologically relevant subtypes of samples.

In the standard setting of differential expression analysis, all genes are tested in two given groups, e.g. disease vs control. In contrast, in our case the groups of samples are undefined and our aim is to discover them. If genes are up-regulated in more than half of all samples, the remaining samples also form a down-regulated module and *vice versa*. Therefore, we are always searching for groups of samples of size not bigger than |*S*|/2. Furthermore, the desired module should not be too small in terms of samples, because a smaller module has a higher probability to appear just by chance. We suggest users to select an appropriate *s*_*min*_ value based on the size of the dataset and intended downstream analysis.

### 2.2 Algorithm

#### 2.2.1 Step 1. Assigning sample sets to edges

In the first step, for each interaction edge *i* connecting genes *u* and *v*, DESMOND identifies a maximal set of samples 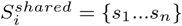 in which both *u* and *v* are differentially expressed compared to 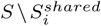. For that, we use a modification of the Rank-Rank Hypergeometric Overlap (RRHO) method (Plaisier *et al.*, 2010) (Figure 1), originally developed for comparison of differential expression profiles obtained in two experiments. It searches for a group of genes significantly enriched in the tops or bottoms of two ranked lists. Basically, this method finds an optimal pair of thresholds, for which the enrichment in tops (bottoms) of the ranked list is the most significant and returns a set of genes with expressions above both thresholds. For two ranked lists of genes, the method creates a 2D-heatmap of the one-sided Fisher’s exact test p-values showing the significance of every pair of threshold values *t*_*u*_, *t*_*v*_, picking a combination corresponding to the most significant overlap.

**Fig. 1.**
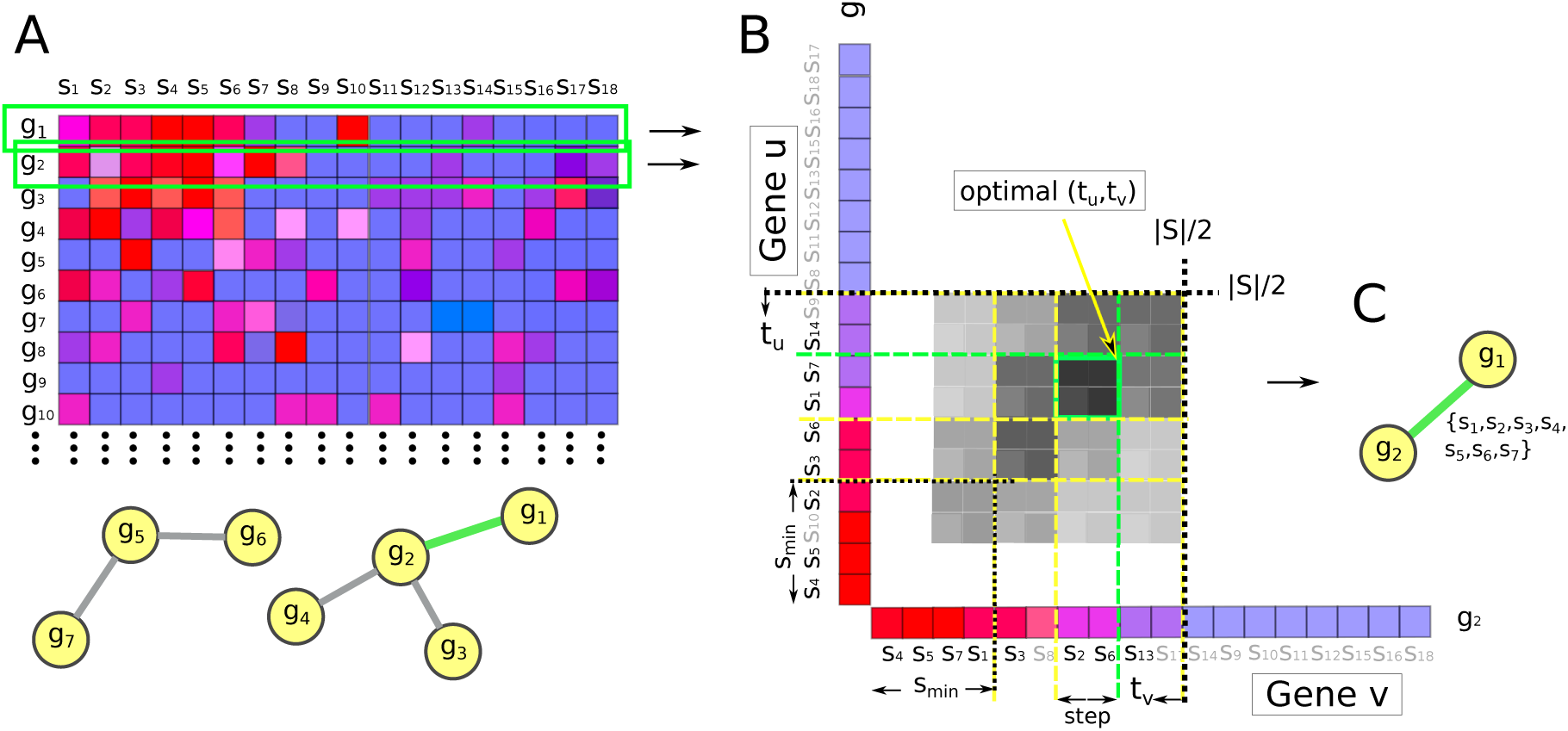
Modified RRHO method used to find the maximal set of samples, in which two interacting genes *g*_1_ and *g*_2_ are up-regulated. A. Input network and expression matrix, red and blue respectively indicate higher and lower expressions. B. Two lists of samples arranged in decreasing order of the expression values of *g*_1_ and *g*_2_. Two thresholds *t*1 and *t*2 move from 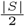 to *s*_*min*_ with step size 2. The intensity of the cell color shows overlap significance for corresponding thresholds. For the case of down-regulation, the same procedure applies, but gene profiles are sorted in ascending order. C. A set of samples *S*^*shared*^ assigned to the edge connecting *g*_1_ and *g*_2_.

For our problem, we modified the RRHO method to find for a given connected pair of genes *u* and *v* a group of samples of size between *s*_*min*_ and 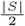, such that both genes are concordantly dysregulated in that sample group. Different from the original method, we move the thresholds from the middle of the lists to the top and stop when achieving the first significant overlap and averaged |*SNR*| value above *SNR*_*min*_. This *SNR*_*min*_ threshold value could be explicitly defined by the user or estimated based on the data (see section 2.2.3 for details). A maximal set of samples 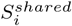 in whose expressions of *u* and *v* are both above the thresholds and whose *avg*.|*SNR*| > *SNR*_*min*_ are assigned to the edge *i*. If no significant overlap bigger than *s*_*min*_ found, the edge is excluded from further consideration.

#### 2.2.2 Step 2. Probabilistic edge clustering

In the first step, every edge is assigned a set of samples in which the pair of genes connected by this edge is up-regulated (or down-regulated). In step two, the algorithm groups edges into connected components, such that each component contains edges with similar sets of samples. We represent the output of the first step as a binary matrix *X* = [*x*_*ji*_]_*n×m*_ for *n* edges and *m* samples, such that *x*_*ji*_ = 1 if sample *i* is assigned to edge *j* and *x*_*ji*_ = 0 otherwise. We propose a constrained Bayesian mixture of Bernoulli distributions for clustering the rows of the matrix into expression modules. The underlying distributions of the mixture model are as follows:

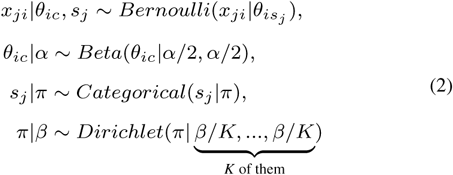

In the above model, the assignments of samples to the edges are modeled as a Bernoulli distribution with parameter *θ*_*ic*_ and a Beta prior, for each sample 1 ≤ *i* ≤ *m* and module 1 ≤ *c* ≤ *K*. The number of modules is set to *K*, equal the number of non-empty edges of the network resulting from step 1. *s*_*j*_, 1 ≤ *s*_*j*_ ≤ *K*, indicates the module to which edge *j* belongs and follows a categorical distribution with parameter *π* and a Dirichlet prior. The model initializes with each edge assigned to a separate module.

We use Gibbs sampling for parameter learning. Each iteration of Gibbs sampling goes over all edges and samples the edge indexes *s*_*j*_ (1 ≤ *j* ≤ *n*). The Gibbs sampling includes two phases: (1) burn-in, consisting of several consecutive iterations for initialization of *s*_*j*_, and (2) sampling, which consists of several iterations throughout which the values of *s*_*j*_ are recorded for further analysis for identification of modules.

At each Gibbs sampling iteration (either in the burn-in or in the sampling phase), to sample the value of *s*_*j*_, we first compute the marginal conditional probability of each *s*_*j*_ belonging to a module *k* as follows:

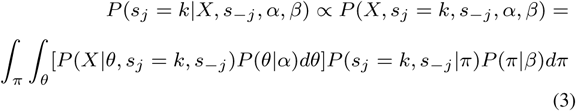

where *s*_−*j*_ indicates the current assignment of all edges except edge *j* to the modules. Because we use conjugate priors (Beta and Dirichlet) the products are in closed form and integrations over *π* and *θ* are straightforward. Keeping the terms that vary with *k*, we get the following conditional probability:

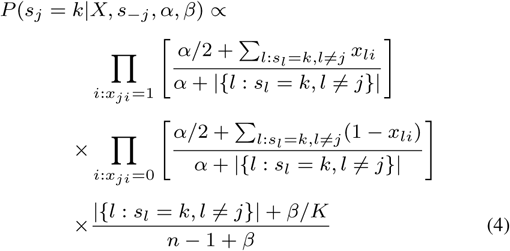

No information is stored about *s*_*j*_ during the burn-in phase. During the sampling phase, which consists of the last 20 iterations before convergence, the values of *s*_*j*_ are recorded. We assume convergence when edge transition probabilities stabilize. Specifically, we compute edge transition probability matrices *P*_*i*_ from the previous 20 model states, starting from *i* + 1-th iteration. Sampling stops when *RMSE*(*Pi*, *P*_*i*+1_) reaches a plateau, more specifically, when the slope of a line fitting RMSE remains between −0.05 and 0.05 during the last five iterations. The final modules are computed as the most frequent value of *s*_*j*_ for each *j* in the last 20 iterations.

#### 2.2.3 Step 3. Post-processing of the resulting modules

In step two, we obtained many overlapping candidate modules containing from zero to many edges. Each module represented a subnetwork, defining a subspace of genes in which samples could be split into two groups differentially expressing these genes. To split all samples into the aforementioned two groups, DESMOND performs 2-means clustering of samples in a subspace of genes representing each module.

Since DESMOND aims to discover subnetworks of differentially expressed genes distinguishing unknown disease subtypes, we exclude all the modules with less than two edges and too low *avg*.|*SNR*|. Users can either explicitly define the *SNR*_*min*_ threshold or draw a certain quantile *q* from the distribution of *avg*.|*SNR*| values computed for 1000 “minimal” biclusters – randomly chosen network edges. Finally, to find more complete gene modules, we merge interconnected modules, dysregulated in the same samples. This is necessary because

- reference biological networks are incomplete (Luck *et al.*, 2017),
- local structure of the network, e.g. changes of network connectivity, may force the method to detect parts of a large bicluster as separate smaller biclusters.

Therefore, we recursively merge modules, starting from the pair with the most significant overlap in samples (Bonferroni-adjusted p-value < 0.05). The merge is only allowed if *avg*.|*SNR*| of the resulting bicluster exceeds *SNR*_*min*_. The procedure is repeated until no merge is possible.

### 2.3 Datasets

#### 2.3.1 Synthetic datasets

We used a similar strategy for synthetic expression data generation, as described in literature (Eren *et al.*, 2012; Padilha and Campello, 2017). For every gene, its expression value was sampled from normal distribution 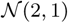 if the gene and sample belonged to a bicluster, or from 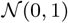 otherwise. Since we had no prior knowledge of the prevailing bicluster sizes in real data, we generated 20 expression matrices and implanted 10 biclusters with the size of 5, 10, 20, 50 or 100 genes and 10, 20, 50 or 100 samples in every matrix. For each bicluster, gene and sample sets were chosen randomly from all genes and samples, i.e. bicluster overlap was allowed.

For each synthetic expression dataset, a scale-free network of 2000 nodes was created using *scale_free_graph* function from Networkx 1.10 python package, implementing the procedure proposed by (Bollobás *et al.*, 2003).

Next, we assigned gene labels to network nodes in a way so that genes from the same bicluster would be connected. First, we assigned to the network genes belonging to exclusive parts of biclusters. For that, we used the approach proposed by (Ghiassian *et al.*, 2015). They have shown that disease-associated genes form compact but not densely connected components on PPI. Therefore, starting from a random node, on every step a neighbor with the highest connectivity p-value (hypergeometric test) is added to the growing component. Next, genes shared by multiple biclusters were assigned to the network, randomly selecting unlabelled nodes connecting the desired biclusters. Finally, background genes were assigned to unlabelled nodes.

#### 2.3.2 Evaluation on synthetic data and the choice of parameters

Ranges of parameter values used in performance evaluation for each method are summarized in Supplementary Table 2. The expected number of biclusters was set to 10 when possible. All other parameters were set to default values. For every combination of parameter values, we applied each method on 20 simulated datasets and calculated Relevance and Recovery scores (Prelić *et al.*, 2006). We considered optimal a combination of parameters corresponding to the highest geometric mean of Relevance and Recovery, averaged over all 20 synthetic datasets. For non-deterministic methods, we computed the average Performance scores in 10 runs.

#### 2.3.3 Evaluation on breast cancer data

We evaluated DESMOND and baseline methods on data collected in two large breast cancer studies, TCGA-BRCA (Liu *et al.*, 2018) and METABRIC (Pereira *et al.*, 2016). In METABRIC cohort, all 1904 gene expression profiles were measured by microarray technology. TCGA-BRCA data comprised of two datasets: 1081 expression profiles were measured by RNA-Seq (TCGA-RNAseq) and 529 by microarrays (TCGA-micro). TCGA-micro and TCGA-RNAseq cohorts were not independent: 517 expression profiles were obtained from the same samples. Since microarray and RNA-seq platforms employ different technologies to estimate gene expression levels, their measurements in the same samples and genes may differ (Robinson *et al.*, 2015). This may affect the results of biclustering, therefore we did not unite these two subsets of expression profiles and applied DESMOND independently on TCGA-RNAseq and TCGA-micro.

Normalized gene expression profiles from TCGA and METABRIC cohorts were downloaded from cBioPortal (Cerami *et al.*, 2012) (http://www.cbioportal.org/). From each cohort, we removed genes expressed in less than 5% of samples, log2-transformed and standardized expressions of remaining genes. Samples from both cohorts were annotated with patient age at diagnosis, stage of the tumor, and molecular subtype. All clinical information was downloaded from cBioPortal and converted into the same format.

We used a human gene network (Huang *et al.*, 2018) derived from the BioGRID (Stark, 2006) network. This network consisted of 258,257 interactions between 16,702 genes. We have chosen BioGRID because it is one of the most comprehensive and frequently updated gene interaction networks for *Homo sapiens*. It comprises curated genetic and protein interactions which are more reliable than computationally predicted interactions. While it provides good coverage of human genes, BioGRID is not too dense. Although most of the edges in this network represent protein interactions, BioGRID still suits our problem because genes with interacting protein products are functionally related.

For evaluation purposes, we kept only 11959 genes presented in all three expression datasets and in the network. We also removed nodes corresponding to genes absent in expression profiles and their adjacent edges from the network before using it (179514 edges remained).

To demonstrate the biological significance of the discovered biclusters, we tested them for associations with Gene Ontology (Consortium, 2016) gene sets and known cancer subtypes using the one-sided exact Fisher’s test. All gene sets used in this work were downloaded from the EnrichR (Kuleshov *et al.*, 2016) website (http://amp.pharm.mssm.edu/Enrichr/).

Overall survival (OS) analysis was performed using the Cox proportional hazards model implemented in Lifelines v0.23.0 (Davidson-Pilon *et al.*, 2019) with age at diagnosis and stage as covariates. All other statistical tests were performed in python using Scipy 1.1.0. Benjamini and Hochberg’s procedure implemented in the gseapy 0.9.9 (Subramanian *et al.*, 2005; Chen *et al.*, 2013) python library was applied for multiple testing correction.

## 3 Results

### 3.1 Synthetic data and parameter tuning

Since previous studies have shown the importance of appropriate parameter setting for method performance (Eren *et al.*, 2012; Sun *et al.*, 2014), each method was run multiple times to find optimal parameter combinations resulting in maximal performance (Supplementary Table 2). Method performances varied widely among tools and different bicluster shapes (Supplementary Figure S1). Interestingly, classic and query-based versions of QUBIC showed similar performances, despite the fact that the latter took into account network information. Although no method outperformed others in all cases, COALESCE demonstrated the best overall performance in this benchmark (on average, 0.63 with default and 0.72 with tuned parameters). DESMOND was the second top-performing method with the average performance of 0.64. It outperformed all other methods for biclusters of sizes 100×100, 50×100 and 5×100 with *α* = 0.5, RRHO p-value threshold *p* = 0.01 and *q* = 0.5. We found that the method was not sensitive to changes of *β*/*K* (Wilcoxon signed-rank test p-values 0.11 and 0.67 for comparison of performances obtained with *β*/*K* set to 1.0 versus 10^4^ and 10^−4^ and other parameters fixed) and therefore set *β*/*K* = 1.0. DESMOND did not perform well on biclusters with a small number of samples. When considering only datasets with biclusters of 20 or more samples, DESMOND on average outperforms all methods including COALESCE (Supplementary Table 3). Therefore, given that DESMOND could not accurately detect the biclusters small in terms of samples, we set *s*_*min*_ to 10% of the whole cohort size in all subsequent experiments. Almost all methods benefited from parameter optimization. DeBi, FABIA, COALESCE, and QUBIC greatly improved their average performances (Figure 2).

**Fig. 2.**
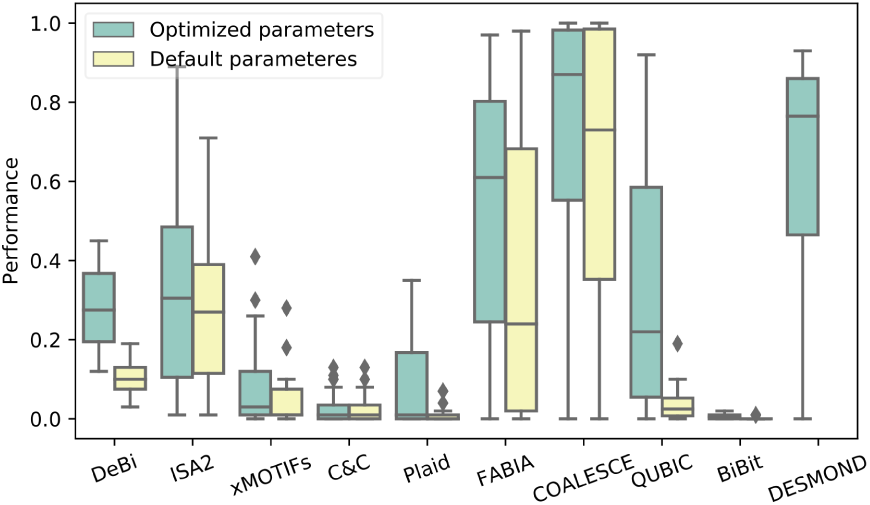
Average performance scores demonstrated by DESMOND and nine baseline methods on 20 synthetic datasets with the default and optimal parameters.

### 3.2 Real data

Five baselines (COALESCE, DeBi, ISA, Fabia, and QUBIC) demonstrated their ability to detect differentially expressed biclusters in synthetic data were chosen and applied on three real-world datasets: TCGA-micro, TCGA-RNAseq, and METABRIC. Each method was run twice: with default parameters and with parameters optimized on synthetic data. Since we were interested in differentially expressed biclusters, we excluded from further analyses all biclusters with *avg*.|*SNR*| < 0.5 (this corresponds to SNR between 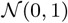 and 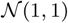) and less than 2 genes or 10 samples.

All methods produced different numbers of biclusters, demonstrating diverse distributions of shapes and *avg*.|*SNR*| values (Figure S2). FABIA run with default parameters identified no biclusters with average |SNR| above 0.5. DeBi did not finish after a week of running with default parameters on the METABRIC dataset and therefore was run on the subset of 500 randomly chosen samples. In contrast, QUBIC identified biclusters with weaker differential expression, when running with optimized parameters than with defaults. Only 8, 2 and 3 biclusters found with optimized parameters in TCGA-micro, TCGA-RNAseq, and METABRIC respectively passed SNR threshold of 0.5. Therefore, below we report the results obtained with optimized parameters for all the methods except QUBIC. Given that the effect of parameter tuning was controversial, we show the results obtained with default and optimized parameters in Supplementary Figures S2-S6.

DESMOND identified 390, 763, and 442 biclusters of 3-157 genes and 53-952 samples in TCGA-micro, TCGA-RNAseq, and METABRIC respectively. Biclusters produced by DESMOND tended to be smaller in terms of genes and bigger in terms of samples than biclusters found by other methods. Compared to the other methods, QUBIC and DESMOND identified biclusters with more pronounced differential expression. In contrast with the synthetic data benchmark, no ground truth was available for real-world breast cancer datasets. Therefore, to evaluate obtained biclusters, we tested the corresponding gene and sample sets for biological significance.

### 3.3 Associations with GO terms

To demonstrate that the identified biclusters are composed of functionally coherent genes, we tested obtained gene sets for overlap with known sets of functionally related genes from Gene Ontology (GO). Most of the biclusters identified by DESMOND were significantly enriched with at least one GO term. Owing to network constraints, the proportion of GO-enriched biclusters was higher for DESMOND than for other methods, including QUBIC which also takes into account the network data (Figure 3).

**Fig. 3.**
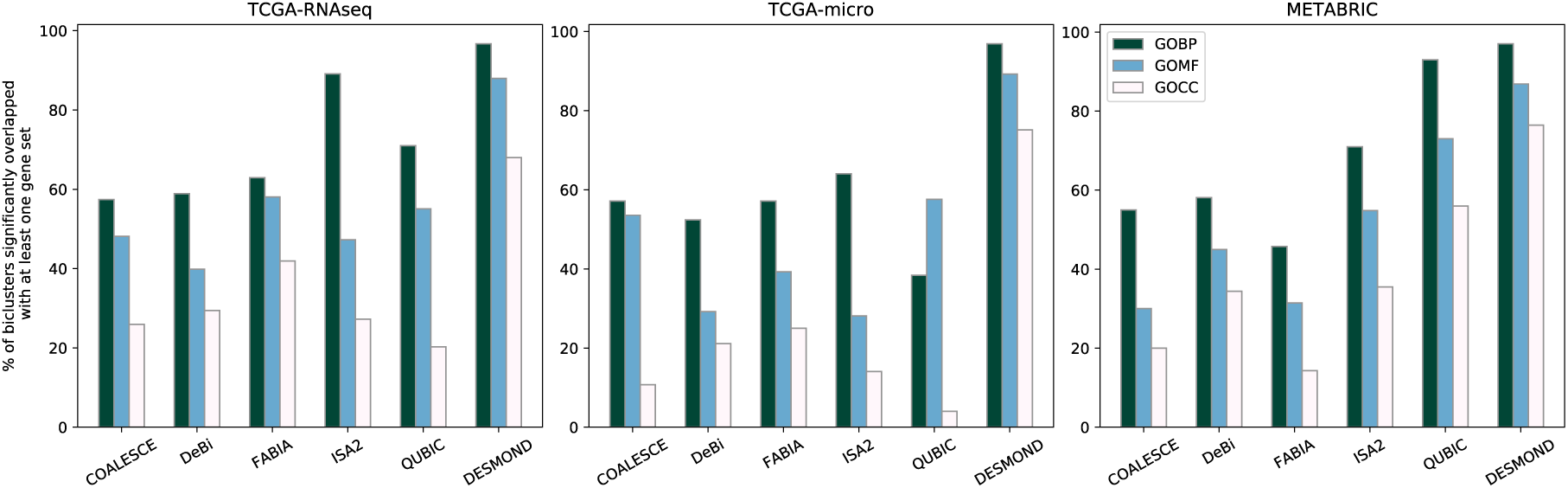
Percentage of gene clusters significantly (BH-adjusted p-value<0.05) overlapping with at least one functionally related gene set from GO Biological Process (GOBP), GO Molecular Function (GOMF) and GO Cellular Component (GOCC). All methods except QUBIC were run with optimized parameters.

To prove that DESMOND performance in this test was superior not only due to network constraints, we generated 100 sets of random subnetworks of the same sizes as DESMOND biclusters. Percent of enriched gene sets was always higher for DESMOND biclusters than for any random set of subnetworks (empirical p-value<0.01). It is important to note that almost all of the random (93-94%) and DESMOND subnetworks (97%) were significantly associated with at least one GOBP term. For GOMF and GOCC the advantage of DESMOND bicluster over random subnetworks was more pronounced (Supplementary Table 4).

### 3.4 Associations with clinical variables

All methods were able to identify many biclusters, significantly (BH-adjusted hypergeometric p-value<0.05) enriched by samples annotated with known breast cancer subtypes. Almost all biclusters found by DESMOND were associated with at least one molecular subtype of breast cancer. Only 0.5-3.3% of DESMOND biclusters showed no significant over-or under-representation of any molecular subtype. In contrast, COALESCE and DeBi produced larger fractions of biclusters not associated with any subtype, up to 68% and 91% of all reported biclusters.

However, although many biclusters were significantly associated with one or several subtypes, only a few of them demonstrated a strong overlap with the associated subtype in terms of the Jaccard similarity. Several methods including DESMOND managed to identify biclusters with strong overlap with Luminal A (LumA) and Basal subtypes (Jaccard similarities about 0.5-0.9, Supplementary Figure S4). ISA2 applied with default parameters identified biclusters relatively strongly (Jaccard similarities about 0.5) overlapping with Her-2 subtype in TCGA. For all other subtypes, overlaps with the most significantly enriched biclusters were weaker.

All identified biclusters were further tested for association with overall survival (OS) using Cox proportional hazards model. DESMOND and DeBi produced more biclusters significantly associated with overall survival compared to other methods (Figures 4, S5, and Supplementary Table 5). Surprisingly, no methods except DeBi with optimized parameters identified OS-associated biclusters in TCGA-micro. However, similarity between OS-associated biclusters found by DeBi on TCGA-micro and TCGA-RNAseq was not high: pairs of biclusters with the strongest overlap in genes never shared more than two samples.

**Fig. 4.**
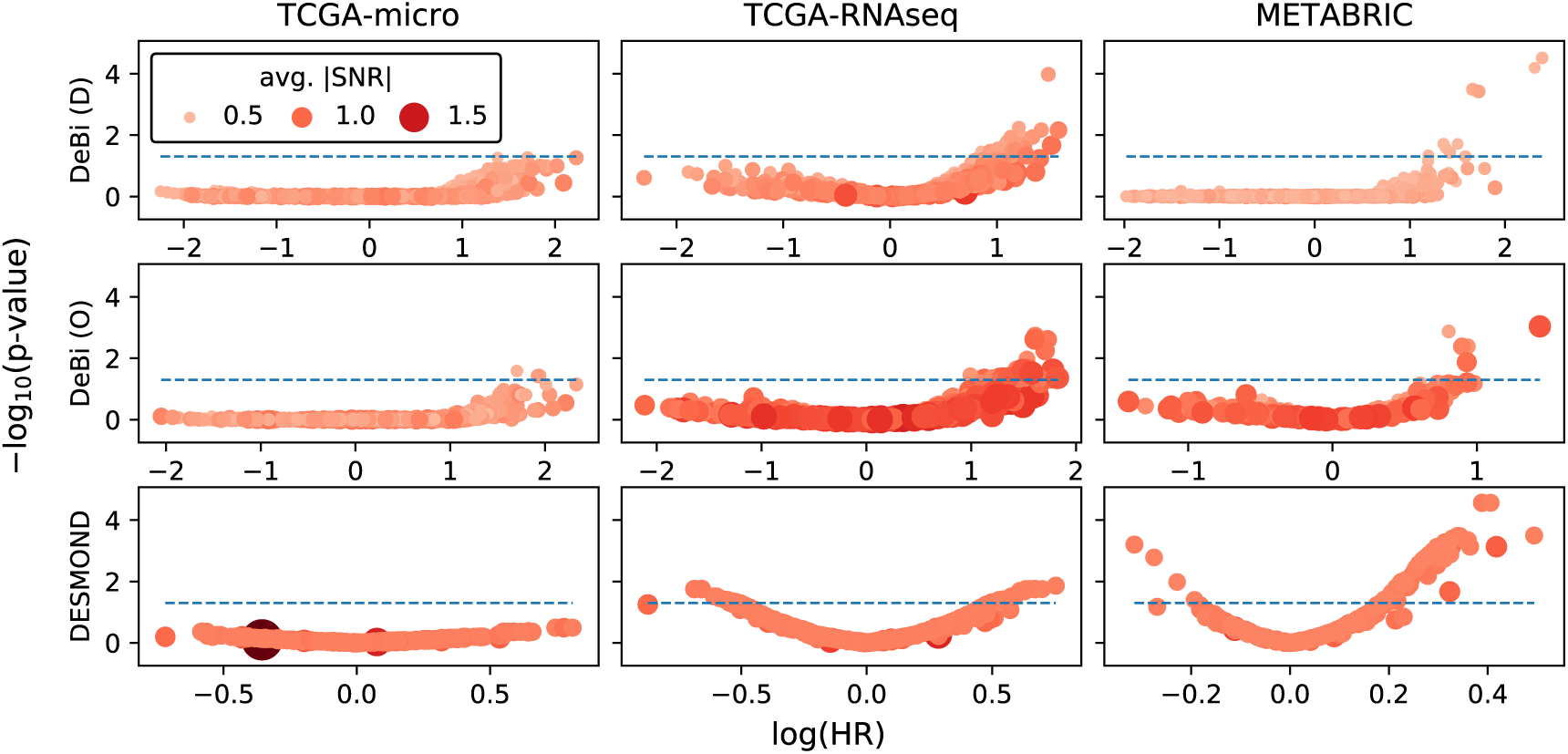
Association of biclusters found by DeBi and DESMOND with overall survival. Every circle represents a bicluster, with size and color intensity proportional to the avg.|SNR|. The X and Y axes show a negative logarithm of adjusted subtype enrichment test p-values and coefficients (logarithm of Hazard Ratio) in Cox regression models fitted for patient sets defined by biclusters. The best biomarkers have higher average SNR and larger positive or negative regression coefficients.

Of all methods, only DESMOND, DeBi and QUBIC identified OS-associated biclusters in both TCGA-RNAseq and METABRIC. QUBIC found only several isolated OS-associated biclusters, overlapping in genes and associated with Basal subtype (Supplementary text). DeBi and DESMOND identified many OS-associated biclusters in these two datasets. Although DeBi identified biclusters with higher HR than DESMOND, the latter produced more similar OS-associated biclusters in TCGA and METABRIC, as we show further.

### 3.5 Reproducibility of found biclusters

To evaluate reproducibility of OS-associated biclusters found by each method in TCGA and METABRIC, we compared best matches between two corresponding sets of biclusters. For each bicluster found in one dataset, its best match in another was determined based on the maximum Jaccard similarity of their gene sets. Since DeBi and QUBIC with optimized parameters produced much larger biclusters in terms of genes than DESMOND, we compared the ratios of observed Jaccard similarities to expected (Figures 5).

**Fig. 5.**
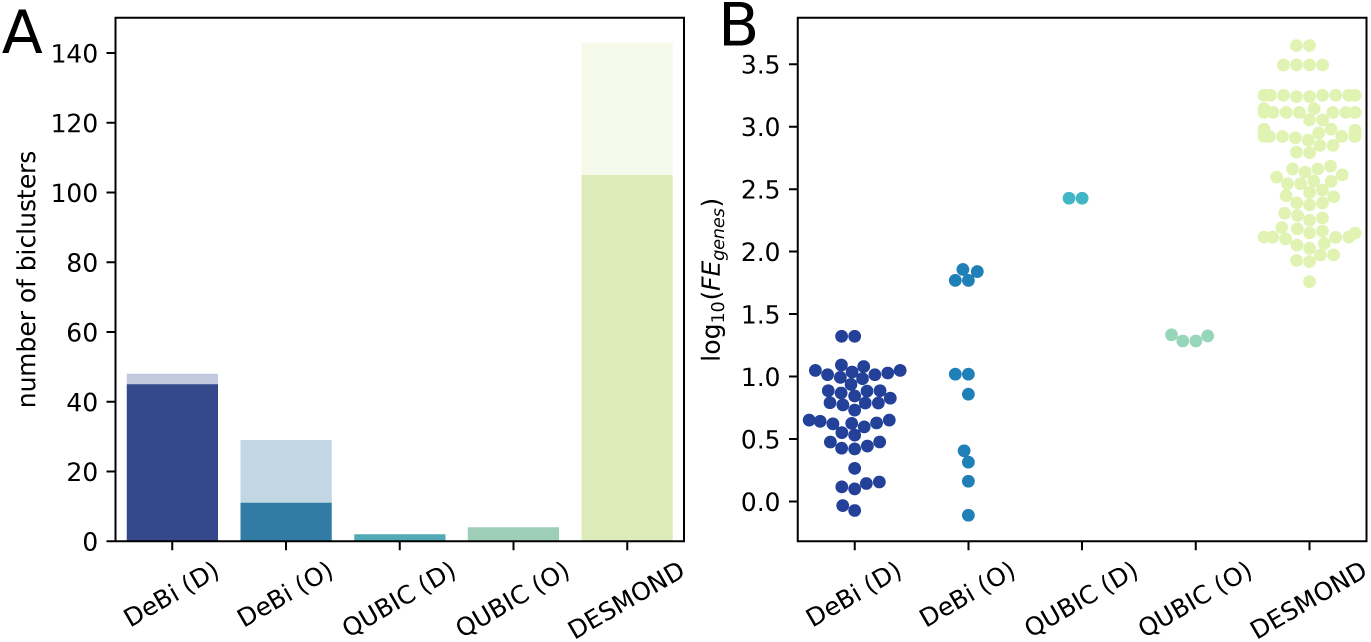
Similarity of OS-associated biclusters found by DeBi, QUBIC and DESMOND in TCGA-RNAseq and METABRIC. A. Total number of OS-associated biclusters found in both datasets. The transparent part of each bar corresponds to biclusters without any match. B. Logarithms of observed Jaccard similarities divided by expected Jaccard similarities.

DeBi and DESMOND identified multiple OS-associated biclusters, but a fraction of their findings had no match between TCGA and METABRIC. All of several biclusters found by QUBIC had a best match strongly overlapping in genes. However, DESMOND biclusters which had a matching partner, tended to demonstrate a higher gain of Jaccard similarity than biclusters produced by the other methods. It is important to note that OS-associated biclusters found by DeBi, QUBIC and DESMOND were not similar in genes and therefore represented different biomarker candidates. The examples of OS-associated biclusters produced by DeBi, QUBIC and DESMOND and composed of similar genes in TCGA and METABRIC are discussed in Supplementary text.

Similarly, we compared all biclusters found by each method in TCGA-BRCA datasets profiled by RNA-seq and microarrays. In this case, both datasets included the same genes and shared 517 samples. Therefore, Jaccard similarity of best matches in genes and samples was calculated considering only shared samples. DESMOND, QUBIC with defaults and FABIA with optimized parameters shown on average a higher gain of Jaccard similarity between best matches than the other methods (Supplementary Figure S6).

## 4 Discussion

In this paper, we presented DESMOND, a new method for the identification of dysregulated gene modules – connected groups of genes up- or down-regulated in unknown subgroups of samples. We applied DESMOND to synthetic and real-world datasets and compared its performance with state-of-the-art biclustering methods.

In the benchmark on synthetic data, DESMOND was the second best performing method, exceeded only by COALESCE. The latter, however, did not outperform the others on real data, apparently, because our synthetic datasets do not reflect the whole complexity of real data. In particular, we did not explicitly model gene co-expressions and used a scale-free network, which may be unrealistic (Broido and Clauset, 2019).

We demonstrated the capability of DESMOND to identify biologically meaningful subsets of genes and samples in real-world breast cancer datasets. However, we note that, similar to some other biclustering methods, DESMOND produced many modules overlapping in their gene sets. This happens because DESMOND clusters interactions between genes, and tends to produce strongly overlapping but different gene clusters from densely connected regions of the network. During the post-processing step, DESMOND merges the strongly overlapping modules to partially address this issue. Further reduction of the redundancy between modules in the first steps of DESMOND remains a direction for future development.

We also note that we did not investigate the effect of different gene networks on the results and tested it only on the BioGRID network. The method may not perform well on a regulatory network, in which co-regulated genes are not connected directly. Also, the network should not be too dense, e.g. like composite functional networks. If dysregulated genes already form a connected component, adding more edges to this component would only increase runtime.

Nevertheless, despite these limitations, we have shown that owing to its ability to consider gene interactions, DESMOND produced more GO-enriched gene clusterings on the breast cancer datasets than other biclustering methods. DESMOND, QUBIC, and DeBi managed to identify different OS-associated biclusters in both breast cancer cohorts. Compared to its competitors, DESMOND tended to identify OS-associated biclusters more similar in genes in independent breast cancer datasets. The better reproducibility of DESMOND biclusters may be explained by the usage of network constraints. Higher stability of the results is desirable for the discovery of gene signatures reproducible in independent studies, regardless of the expression profiling method used. Finding of reproducible OS-associated biclusters may point to the presence of the internal heterogeneity beyond established molecular subtypes and requires further investigation.

## Acknowledgements

We would like to thank Ralf Hofestädt and Maren Kleine (Bielefeld University), Felix Frenkel, Ravshan Ataullakhanov, Aleksander Bagaev, Nikita Kotlov and colleagues from Boston Gene LLC, Nikolay Zolotarev (MPI of Immunobiology and Epigenetics), and four anonymous reviewers of ISMB’19 for their insightful comments and suggestions.

## Funding

This study was supported by the International Deutsche Forschungsgemeinschaft Research Training Group GRK 1906 and the AG Bioinformatik and Medical Informatik of Bielefeld University.

## Notes

https://github.com/ozolotareva/DESMOND

### Competing Interest Statement

The authors have declared no competing interest.

## References

Bergmann, S., Ihmels, J. and Barkai, N. (2003). Iterative signature algorithm for the analysis of large-scale gene expression data. Physical Review E, 67(3).

Bollobás, B., Borgs, C., Chayes, J. et al.(2003). Directed scale-free graphs. In Proceedings of the Fourteenth Annual ACM-SIAM Symposium on Discrete Algorithms, SODA’03, pages 132–139, Philadelphia, PA, USA. Society for Industrial and Applied Mathematics.

Broido, A.D. and Clauset, A. (2019). Scale-free networks are rare. Nature Communications, 10(1).

Cerami, E., Gao, J., Dogrusoz, U. et al.(2012). The cBio cancer genomics portal: An open platform for exploring multidimensional cancer genomics data. Cancer Discovery, 2(5), 401–404.

Chen, E.Y., Tan, C.M., Kou, Y. et al.(2013). Enrichr: interactive and collaborative HTML5 gene list enrichment analysis tool. BMC Bioinformatics, 14(1), 128.

Cheng, Y. and Church, G.M. (2000). Biclustering of expression data. In Proceedings of the Eighth International Conference on Intelligent Systems for Molecular Biology, pages 93–103. AAAI Press.

Chowdhury, S.A. and Koyutürk, M. (2009). IDENTIFICATION OF COORDINATELY DYSREGULATED SUBNETWORKS IN COMPLEX PHENOTYPES. In Biocomputing 2010, pages 133–144. WORLD SCIENTIFIC.

Consortium, G.O. (2016). Expansion of the gene ontology knowledgebase and resources. Nucleic Acids Research, 45(D1), D331–D338.

Dao, P., Wang, K., Collins, C. et al.(2011). Optimally discriminative subnetwork markers predict response to chemotherapy. Bioinformatics, 27(13), i205–i213.

Davidson-Pilon, C., Kalderstam, J., Zivich, P. et al.(2019). Camdavidsonpilon/lifelines: v0.23.0.

Eren, K., Deveci, M., Kucuktunc, O. et al.(2012). A comparative analysis of biclustering algorithms for gene expression data. Briefings in Bioinformatics, 14(3), 279–292.

for Research on Cancer, I.A. (2012). WHO Classification of Tumours of the Breast (Medicine). World Health Organization.

Ghiassian, S.D., Menche, J. and Barabási, A.L. (2015). A DIseAse MOdule detection (DIAMOnD) algorithm derived from a systematic analysis of connectivity patterns of disease proteins in the human interactome. PLOS Computational Biology, 11(4), e1004120.

Gradishar, W.J., Anderson, B.O., Balassanian, R. et al.(2017). NCCN guidelines insights: Breast cancer, version 1.2017. Journal of the National Comprehensive Cancer Network, 15(4), 433–451.

He, Z. and Zhou, J. (2008). Empirical evaluation of a new method for calculating signal-to-noise ratio for microarray data analysis. Applied and environmental microbiology, 74 10, 2957–66.

Hochreiter, S., Bodenhofer, U., Heusel, M. et al.(2010). FABIA: factor analysis for bicluster acquisition. Bioinformatics, 26(12), 1520–1527.

Huang, J.K., Carlin, D.E., Yu, M.K. et al.(2018). Systematic evaluation of molecular networks for discovery of disease genes. Cell Systems, 6(4), 484–495.e5.

Huttenhower, C., Mutungu, K.T., Indik, N. et al.(2009). Detailing regulatory networks through large scale data integration. Bioinformatics, 25(24), 3267–3274.

Ideker, T., Ozier, O., Schwikowski, B. et al.(2002). Discovering regulatory and signalling circuits in molecular interaction networks. Bioinformatics, 18(Suppl 1), S233–S240.

Khakabimamaghani, S. and Ester, M. (2015). BAYESIAN BICLUSTERING FOR PATIENT STRATIFICATION. In Biocomputing 2016. WORLD SCIENTIFIC.

Kuleshov, M.V., Jones, M.R., Rouillard, A.D. et al.(2016). Enrichr: a comprehensive gene set enrichment analysis web server 2016 update. Nucleic Acids Research, 44(W1), W90–W97.

Lazzeroni, L. and Owen, A. (2000). Plaid models for gene expression data. Stat Sin., 12, 61–86.

Li, G., Ma, Q., Tang, H. et al.(2009). QUBIC: a qualitative biclustering algorithm for analyses of gene expression data. Nucleic Acids Research, 37(15), e101–e101.

Liu, J., Lichtenberg, T., Hoadley, K.A. et al.(2018). An integrated TCGA pan-cancer clinical data resource to drive high-quality survival outcome analytics. Cell, 173(2), 400–416.e11.

Love, M.I., Huber, W. and Anders, S. (2014). Moderated estimation of fold change and dispersion for RNA-seq data with DESeq2. Genome Biology, 15(12).

Luck, K., Sheynkman, G.M., Zhang, I. et al.(2017). Proteome-scale human interactomics. Trends in Biochemical Sciences, 42(5), 342–354.

McCarthy, D.J., Chen, Y. and Smyth, G.K. (2012). Differential expression analysis of multifactor RNA-seq experiments with respect to biological variation. Nucleic Acids Research, 40(10), 4288–4297.

McClellan, J. and King, M.C. (2010). Genetic heterogeneity in human disease. Cell, 141(2), 210–217.

Mishra, D. and Sahu, B. (2011). A signal-to-noise classification model for identification of differentially expressed genes from gene expression data. In 2011 3rd International Conference on Electronics Computer Technology. IEEE.

Mitra, K., Carvunis, A.R., Ramesh, S.K. et al.(2013). Integrative approaches for finding modular structure in biological networks. Nature Reviews Genetics, 14(10), 719–732.

Murali, T. and Kasif, S. (2003). Extracting conserved gene expression motifs from gene expression data. Pacific Symposium of Biocomputing, pages 77–88.

Padilha, V.A. and Campello, R.J.G.B. (2017). A systematic comparative evaluation of biclustering techniques. BMC Bioinformatics, 18(1).

Pereira, B., Chin, S.F., Rueda, O.M. et al.(2016). The somatic mutation profiles of 2, 433 breast cancers refine their genomic and transcriptomic landscapes. Nature Communications, 7(1).

Perou, C.M., Sørlie, T., Eisen, M.B. et al.(2000). Molecular portraits of human breast tumours. Nature, 406(6797), 747–752.

Plaisier, S.B., Taschereau, R., Wong, J.A. et al.(2010). Rank–rank hypergeometric overlap: identification of statistically significant overlap between gene-expression signatures. Nucleic Acids Research, 38(17), e169–e169.

Pontes, B., Giráldez, R. and Aguilar-Ruiz, J.S., (2015). Biclustering on expression data: A review. Journal of Biomedical Informatics, 57, 163–180.

Prelić, A., Bleuler, S., Zimmermann, P. et al.(2006). A systematic comparison and evaluation of biclustering methods for gene expression data. Bioinformatics, 22(9), 1122–1129.

Reiss, D.J., Plaisier, C.L., Wu, W.J. et al.(2015). cMonkey2: Automated, systematic, integrated detection of co-regulated gene modules for any organism. Nucleic Acids Research, 43(13), e87–e87.

Ritchie, M.E., Phipson, B., Wu, D. et al.(2015). limma powers differential expression analyses for RNA-sequencing and microarray studies. Nucleic Acids Research, 43(7), e47–e47.

Robinson, D.G., Wang, J.Y. and Storey, J.D. (2015). A nested parallel experiment demonstrates differences in intensity-dependence between RNA-seq and microarrays. Nucleic Acids Research, page gkv636.

Rodriguez-Baena, D.S.,, Perez-Pulido, A.J., and Aguilar-Ruiz, J.S., (2011). A biclustering algorithm for extracting bit-patterns from binary datasets. Bioinformatics, 27(19), 2738–2745.

Saelens, W., Cannoodt, R. and Saeys, Y. (2018). A comprehensive evaluation of module detection methods for gene expression data. Nature Communications, 9(1).

Serin, A. and Vingron, M. (2011). DeBi: Discovering differentially expressed biclusters using a frequent itemset approach. Algorithms for Molecular Biology, 6(1).

Stark, C. (2006). BioGRID: a general repository for interaction datasets. Nucleic Acids Research, 34(90001), D535–D539.

Subramanian, A., Tamayo, P., Mootha, V.K. et al.(2005). Gene set enrichment analysis: A knowledge-based approach for interpreting genome-wide expression profiles. Proceedings of the National Academy of Sciences, 102(43), 15545–15550.

Sun, P., Speicher, N.K., Röttger, R. et al.(2014). Bi-force: large-scale bicluster editing and its application to gene expression data biclustering. Nucleic Acids Research, 42(9), e78–e78.

Sweeney, T.E., Haynes, W.A., Vallania, F. et al.(2016). Methods to increase reproducibility in differential gene expression via meta-analysis. Nucleic Acids Research, 45(1), e1–e1.

Turner, H., Bailey, T. and Krzanowski, W. (2005). Improved biclustering of microarray data demonstrated through systematic performance tests. Computational Statistics & Data Analysis, 48(2), 235–254.

Xie, J., Ma, A., Fennell, A. et al.(2018). It is time to apply biclustering: a comprehensive review of biclustering applications in biological and biomedical data. Briefings in Bioinformatics.

Zhang, M., Yao, C., Guo, Z. et al.(2008). Apparently low reproducibility of true differential expression discoveries in microarray studies. Bioinformatics, 24(18), 2057–2063.

